# Cortical inhibitory network selects cerebellar signals for movement initiation

**DOI:** 10.1101/2020.10.20.346775

**Authors:** Abdulraheem Nashef, Oren Cohen, Steve I. Perlmutter, Yifat Prut

**Affiliations:** Department of Medical Neurobiology, IMRIC and ELSC, The Hebrew University, Hadassah Medical School, Jerusalem 9112102, Israel; Department of Physiology & Biophysics and the Washington National Primate Research Center, Box 357330, University of Washington, Seattle, Washington 98195, USA

**Keywords:** Thalamocortical, Motor control, Feed-forward inhibition, Iontophoresis, PV+ interneurons, Cerebellum, Non-human primates

## Abstract

The onset of voluntary movements is driven by coordinated firing across a large population of motor cortical neurons. This pattern of activity is determined by both local interactions and long-range corticocortical and subcortical inputs. The way remote areas of the brain communicate to effectively drive movement is still unclear. We addressed this question by studying an important pathway through which the cerebellum communicates, via the motor thalamus, with the motor cortex. We found that similar to the sensory cortices, thalamic input to the motor cortex triggers feedforward inhibition by directly contacting inhibitory cells via particularly effective GluR2- lacking AMPA receptors blocked by NASPM. Based on these results, we constructed a classifier for SCP-responsive cortical cells to identify pyramidal and PV interneurons and study their role in controlling movements. The findings indicate that PV and pyramidal cells are co-driven by TC input in response to activation of the CTC pathway. During task performance, PV and pyramidal cells had comparable relations to movement parameters (directional tuning and movement duration). However, PV interneurons exhibited stronger movement-related activity that preceded the firing of pyramidal cells. This seemingly counterintuitive sequence of events where inhibitory cells are recruited more strongly and before excitatory cells may in fact enhance the signal-to-noise ratio of cerebellar signals by suppressing other inputs and prioritizing the excitatory synchronized volley from the TC system which occurs at the right time to overcome the inhibitory signal. In this manner, the CTC system can shape cortical activity in a way that exceeds its sheer synaptic efficacy.

## INTRODUCTION

Motor commands for voluntary movements result from the integration of information from multiple cortical and subcortical sources by the motor cortex (Middleton and Strick, 2000). Numerous studies have explored the role of local motor cortical dynamics in producing actions (Churchland et al., 2012; Kaufman et al., 2014; Kaufman et al., 2013; Shenoy et al., 2013). However, the way remote brain regions communicate with the motor cortex and take an effective part in driving movements is still unclear. To investigate this question, we focused on the cerebellar-thalamocortical (CTC) pathway, through which the cerebellum influences motor cortical activity. Cerebellar signals participate in shaping motor cortical commands and play a specific role in the regulation of motor timing coordination (Bastian et al., 1996; Holmes, 1917, 1939; Ivry and Keele, 1989). This means that despite the large distance cerebellar information needs to travel, and the limited number of synapses onto motor cortical neurons (Bopp et al., 2017; Schoonover et al., 2014) this pathway exerts online impact on motor output. In sensory (Cruikshank et al., 2007; Gabernet et al., 2005) and frontal (Delevich et al., 2015) cortical areas, thalamocortical inputs produce feedforward inhibition (FFI) of cortical activity, which dictates a temporal window for information processing (Pouille and Scanziani, 2001). In the motor system, there is indirect evidence for a cerebellar role in braking and inhibiting motor cortical circuitry (Hore and Flament, 1988; Nashef et al., 2018a) and unwarranted movements (Babinski, 1913). We hypothesized that a feedforward mechanism in the CTC system may mediate and prioritize the role of cerebellar signals in timing the onset of voluntary movements.

To test this hypothesis, we combined stimulation of the CTC pathway, electrophysiological recordings of motor cortical neurons, and pharmacological perturbations of transmission through this pathway in behaving primates. This enabled us to positively identify GABAergic interneurons and pyramidal neurons that integrate CTC information in the motor cortex. Based on studies in the sensory system of rodents, where the circuitry of the FFI relies on particularly powerful synapses by TC fibers made selectively onto fast-spiking parvalbumin positive (PV) neurons (Bruno, 2011; Cruikshank et al., 2007; Ji et al., 2015; Yu et al., 2019), we suggest that these interneurons are in fact PV cells. The properties of this subset of cells were used to construct a classifier for PV and pyramidal neurons out of all the cells responsive to stimulation of the superior cerebellar peduncle (SCP). During task performance, the movement-related activity of PV neurons preceded that of pyramidal cells and their directional tuning was slightly broader. In addition, the activity of PV cells was strongly correlated with the kinematic properties of the movements. Finally, the firing of PV and pyramidal cells was specifically correlated around movement onset, indicating strong and specific cross-interactions between these cell groups.

Our results suggest that the CTC system relies on FFI in the motor cortex in a way similar to the circuitry found in cortical sensory areas of rodents. The functional consequences of this circuitry are the strong coupling between SCP-responsive PV and pyramidal cells and the task-related activity of PV cells preceding that of the pyramidal cells. We posit that this counterintuitive sequence of events serves to assign a high priority to cerebellar signals over other competing inputs. In this manner PV neurons play an important role in steering cortical activity at movement onset to efficiently translate cerebellar input into timing signals for motor commands. Finally, the inhibitory control exerted by the cerebellum on motor output maybe mediated, in part, by the specific potent recruitment of motor cortical interneurons.

## RESULTS

Our previous findings indicated that cerebellar signals routed through the CTC system efficiently recruit an extensive population of cortical cells throughout the motor cortex (Nashef et al., 2018a). Here we aimed to dissect the local cortical circuitry that integrates these signals to elucidate the involvement of different subclasses of cortical neurons in controlling the timing of movements. To do so, we took advantage of the organizational principles of the TC system in the sensory and frontal areas of rodents where TC axons make distinctive connections with excitatory and inhibitory cells (Cruikshank et al., 2007; Gabernet et al., 2005). We implanted stimulating electrodes in the SCP (Figure 1) of monkeys (n=3) that were trained to wear an exoskeleton (Kinarm system) and perform a center-out reaching task (Figure 1A-B). The behavioral task used here was designed to encourage the monkeys to move as fast as possible by predicting the time of the “go” signal (Nashef et al., 2019). Neural activity was recorded from arm-related motor cortical areas using multiple single electrodes in 3 monkeys (Figure 1C); in two of these animals, neurons were also tested using a 3-barrel recording pipette (Thiele et al., 2006). This experimental approach (Figure 1D) enabled us to record the activity of motor cortical cells during movement, identify their response to CTC activation, and in some cases pharmacologically perturb their activity by iontophoretic application of neuroactive compounds.

**Figure 1:**
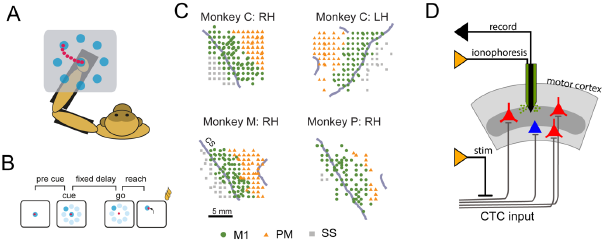
Experimental Design. (A) Schematic illustration of the experimental setup. Monkeys were trained to wear an exoskeleton and control a cursor that appeared on a horizontally positioned screen. The movement of the monkeys was constrained to planar, shoulder-elbow reaching movements. (B) Behavioral paradigm. The sequence of events composing a single trial included a pre-cue period, target onset (where 1 of 8 equally distributed targets appeared), a delay period, and a go signal, after which the monkey had to acquire the target within a predefined movement time. Correct performance resulted in a reward (a drop of applesauce). (C) Cortical maps of recorded sites obtained from monkey C (right hemisphere, RH; left hemisphere, LH) and monkeys M and P (right hemisphere only). CS, central sulcus. Scale bar: 5 mm. Green circles: sites in the primary motor cortex (M1). Orange triangles: sites in premotor cortex (PM). (D) Stimulating electrode was chronically implanted in the cerebellar-thalamo-cortical (CTC). Three-barrel pipette with a tungsten core was used to record and apply drugs iontophoretically in the motor cortex while monkeys performed the task.

### Identifying motor cortical neurons that integrate cerebellar signals

The connectivity pattern between TC fibers and specific classes of neurons has a major impact on the way subcortical input affects cortical activity. In sensory areas, TC inputs contact both pyramidal and PV cells, thus generating a feedforward inhibitory connection. In fact, the contacts between TC axons and cortical inhibitory interneurons are almost exclusively confined to PV cells (Ji et al., 2015), and these synapses are mediated by a specific type of AMPA receptor which lacks the GluR2 subunit (Hull et al., 2009). These receptors are unique in their calcium permeability, their particularly large synaptic currents, and the specificity of the blocker 1-naphythyl acetyl spermine or NASPM (Hull et al., 2009). Importantly, GluR2-lacking AMPA receptors are not found on pyramidal cells of the adult brain (Geiger et al., 1995; Kumar et al., 2002; McBain and Dingledine, 1993).

Based on these data we employed a two-step identification criterion for PV neurons in the CTC pathway (Figure 2A). We identified cells (n=360) that were part of the CTC system by their early excitatory response to single pulse SCP stimulation (Nashef et al., 2018a). Out of this group, we tested 58 neurons for changes in the SCP-triggered response after blocking GluR-2 receptors by iontophoretically applying NASPM from the barrel of the recording pipette. Cells that decreased their early excitatory response to SCP stimulation in the presence of NASPM were identified as PV interneurons whereas cells that either maintained or increased their evoked response were defined as pyramidal cells (Figure 2B). Of the 58 tested cells, 28% (n=16) decreased their response in the presence of NASPM, and were classified as PV interneurons. The remaining 42 cells (72%) maintained (or increased) their response to stimulation, and were classified as pyramidal cells (Pyr). In the presence of NASPM, the SCP-triggered response of PV interneurons exhibited a significant decrease in response amplitude (Figure 2C), pre-stimulus firing rate, area under the excitatory peak of the response curve, and area of the inhibitory phase of the response (Wilcoxon’s signed-rank test; p<0.001 for all parameters). In contrast to these robust changes, the response pattern of pyramidal neurons to SCP stimulation was unaffected (Figure 2C). In addition, neurons classified as PV cells had significantly narrower action potential (AP) widths (measured as the time from trough to peak) than pyramidal neurons (Pyr=0.46±0.02 ms; PV=0.35±0.03 ms; Wilcoxon’s rank sum test, p<0.005). These findings thus indicate that in the CTC system of primates, TC fibers connect directly to both excitatory (pyramidal) and inhibitory (presumably PV) cells in a scheme comparable to the FFI reported for sensory systems in rodents.

**Figure 2:**
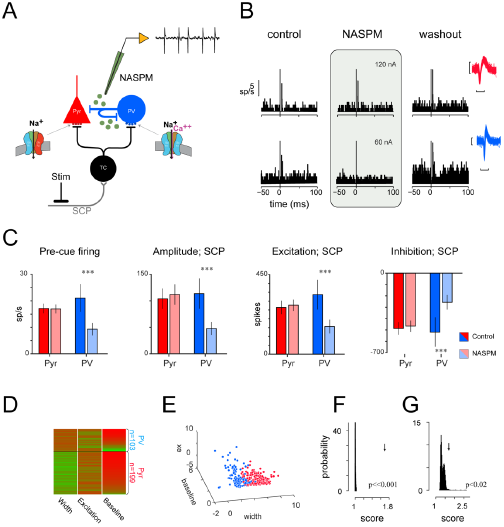
Pyramidal and Parvalbumin neurons identification. (A) Experimental design for iontophoresis testing. Ascending axons of thalamocortical (TC) neurons make contact with parvalbumin (PV) and pyramidal (Pyr) neurons, the synapse between the TC fibers and PV neurons is mediated by GluR2-lacking AMPA receptors which are Ca^+2^ permeable. A triple-barrel pipette was filled with 1-Naphthyl acetyl spermine trihydrochloride (NASPM) which is a specific blocker of GluR2-lacking AMPA receptors. During recordings we stimulated the CTC via a stimulation electrode that was chronically implanted in the superior cerebellar peduncle (SCP). We tested the synaptic contact by studying the modulation of response to stimulation during iontophoresis of NASPM. (B) Example of two cells that responded to SCP stimulation and were tested using NASPM. The response is measured using the peri-stimulus time histogram aligned on stimulation onset time (t=0). Response to SCP stimulation was measured, from left to right, during control trials (left column), during iontophoresis of NASPM (shaded column) and during washout trials (right column). In the presence of NASPM one of the cells (upper panel) maintained responding to SCP stimulation but the second cell (lower panel) stopped responding. Based on these properties we defined the cell on the upper row as a Pyramidal cell and the cell on the lower row as a PV cell. The action potential waveforms are shown as well (right) for the two cells by super-positioning 50 randomly selected waveforms emitted by the Pyramidal (red) and PV (blue) cells. (Response: scale bar- 5 sp/s; Waveform: vertical scale bar- 50 μV, horizontal scale bar- 1 ms). (C) Differences in firing properties between Pyramidal (red, n=42) and PV (blue, n=16) cells in control trials (dark hues) and in the presence of NASPM (light hues). From left to right: Baseline of activity when no stimulation was applied (Pyr: 17.1±1.8 to 16.9±1.7; PV: 21.1±3.6 to 9.3±2.4 sp/s); Amplitude of response to SCP stimulation (mean±s.e.m.; control to NASPM; Pyr: 103.8±19.1 to 111.8±19.4; PV: 113.6±23.1 to 47.7±16.1 sp/s); Area under the curve of the excitation period of the SCP response (Pyr: 264.8±38.1 to 276.6±31.8; PV: 336.5±57.7 to 156.1±42.4 spikes); Area under the curve of the inhibition period of the SCP response (Pyr: -487±53.9 to -466.2±49.4; PV: -518±58.3 to -256.7±44.3 spikes). ***p<0.001; Wilcoxon’s sign-rank. (D) Classification of entire population of SCP-responsive cells (n=302). Three features that we considered for cell classification (Width: width of waveform computed from trough to peak; Excitation: area under the early positive part of the SCP-evoked response; Baseline- background activity when no stimulation was applied). The color scale runs from red (low values) to green (high values) and reflects the differences between the two cell classes. Black horizontal line distinguishes between the Pyramidal (bottom; n=199 cells) and PV (top; n=103 cells) populations. (E) A 3-dimensional scatter plot of the three features used for classification. Pyramidal cells are shown as red circles, and PV cells as blue circles. An RCA algorithm (Bar-Hillel et al., 2003; Shental et al., 2002) was used here to optimize the differences between the cell classes. (F) Comparison between mean clustering score obtained for the data (arrow; score=1.68) and the distribution of values obtained for shuffled data (bars; 1.01±0.0003; p<<0.001). The score was calculated as the distance of each point to the center of the other cluster over its distance to the center of its corresponding cluster. (E) Comparison between the mean clustering score calculated for the data (arrow; score=1.68) and the values obtained when using a Gaussian clustering method on 80% of the neurons (n=242 neurons) without using the pre-training population of neurons (identified using iontophoresis) as was done here (bars; 1.4±0.001; p=0.012). See also Figure S1.

In the second step of the identification process, the responses of the cortical cells to SCP stimulation, their background firing rate and AP width were used to construct a mixed classifier (naïve Bayesian and support-vector machine-SVM (Cortes and Vapnik, 1995)) which was applied to the remaining dataset of 302 SCP-responsive cells (Figure 2D). The most clear-cut feature for classification (beyond the actual response to stimulation, which was set as an initial condition) was the width of the waveform, which alone achieved 70% classification accuracy. The pre-stimulus firing level and the magnitude of excitation triggered by SCP stimulation (area under the response curve) improved the classification accuracy to 76% (Pyr: 76.2%; PV: 75%). Although the pre-stimulus firing level and the magnitude of excitation were not significantly different for the training population between the two classes, the three features used here showed a significant multivariate relationship with cell class (multivariate analysis of variance, p<0.005; degrees of freedom=57). Overall, we classified 199/302 (66%) SCP-responsive cortical cells as pyramidal cells and the remaining 103 (34%) as PV cells (Figure 2E). For each neuron, we calculated the ratio of the distance between the neuron’s location in the clustering space (Figure 2E) to the center of the other cluster over the distance to the neuron’s allocated cluster center. We then averaged the ratios for all neurons to define a clustering score as a measure of the goodness of clustering. This score was compared to the distribution of scores obtained in a similar manner on datasets where cluster allocation was random, while keeping the counts for each cell class fixed. The actual score (computed for the real dataset) was located far from the distribution of values calculated for shuffled datasets (Figure 2F, p << 0.001), supporting the classification approach used here. In another test, we compared the results of classification from the training set to a “blind” classification procedure using a Gaussian mixture distribution classifier of two subpopulations. We found that the clustering based on the training dataset yielded higher clustering scores than the blind classification (p<0.02; Figure 2G). Thus, the training-based classifier used here outperformed untrained clustering methods used in the past (Katai et al., 2010; Trainito et al., 2019).

### Movement-related activity of PV interneurons precedes activity of pyramidal cells

We compared the movement-related activity of identified cells during task performance. In our dataset, inhibitory cells had a stronger and more reliable response to SCP stimulation (Figure S1). This is consistent with previous studies showing that the cortical response to activation of the CTC system is dominated by inhibition (Hore and Flament, 1988; Nashef et al., 2018a). This differential recruitment of specific cell types by CTC signals was expected to promote distinct patterns of activity in these cells during movement. To test this hypothesis, we compared the response pattern of pairs of PV and Pyr cells recorded simultaneously on one or two electrodes. Simultaneous recordings eliminate the possibility of confounds due to session-to-session variability in motor behavior. We found significantly different movement-related properties in PV and Pyr neurons (Figure 3). PV cells had a higher background firing rate (Figure 3A, Wilcoxon’s signed-rank test; p<0.05), and a higher task-related response amplitude around movement onset time (Figure 3B; p<0.003). However, the most remarkable difference between the two cell types was the tendency of PV cells to increase their movement-related firing earlier than Pyr cells. Figure 3C shows an example of movement-related activity of a PV cell and a Pyr cell recorded simultaneously. In this example, the onset of the increased movement-related activity of the PV cell preceded that of the Pyr cell by 360 ms and the peak firing was 4.7 sp/s higher. The propensity of PV cells for an earlier and larger amplitude response was also present at the population level (Figure 3D). To compensate for the higher activity of PV compared to Pyr cells, we normalized the firing rate of each neuron to its peak activity (which was thus defined as 100%) and identified the time at which activity reached 20% of the maximal peak value (Figure 3E). We found that PV neurons reached this threshold about 200 ms before pyramidal cells.

**Figure 3:**
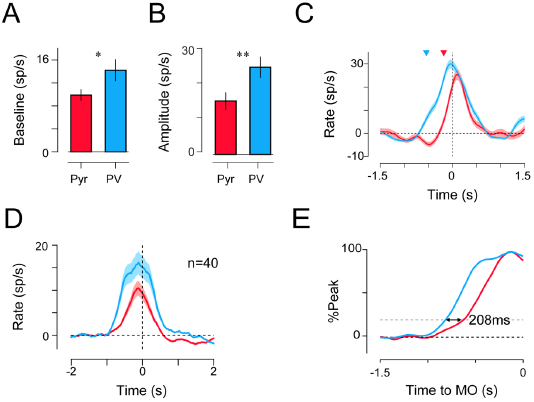
Task-related activity of simultaneously recorded PV and pyramidal cells during reaching. (A) Baseline activity computed for simultaneously recorded Pyramidal (red) and PV (blue) cells during a 1 second time window spanning from -2 to -1 s relative to movement onset, when no movement took place (n=40 pairs; Pyr: 9.9±1.02; PV: 14.2±1.9 sp/s; Wilcoxon’s sign rank: p<0.05). (B) Response amplitude measured as the maximal firing rate around movement onset for the same group of cells as in A. (Pyr: 25.3±2.8; PV: 39.4±3.8 sp/s; p<0.003). (C) Example of task-related activity around movement onset for Pyr (red) and PV (blue) cells that were recorded simultaneously. Shaded area depicts the standard error of the mean and arrowheads highlight the response onset time calculated for each neuron (Pyr onset=-180 ms; PV onset=-540 ms). (D) Average (± s.e.m) task-related activity of simultaneously recorded pyramidal and PV cells. Activity is aligned on movement onset and baseline activity was subtracted. (E) Normalized neural activity obtained by dividing by peak firing rate around movement onset and expressed in percentages. Difference between average profiles was measured to be 208 ms which was statistically significant (Wilcoxon’s sign-rank; p<0.05 n= 40 cell pairs).

### Encoding of motor parameters by different cell classes

Next, we tested whether the correlation between movement-related signals and specific kinematic parameters was different for the two cell classes. We found that the tuning of PV and Pyr cells for the direction of arm movement had similar widths (p < 0.23). The same was true for other measures for tuning selectivity, such as the half-width at half height (HWHH) and the orientation selectivity (measured as the 1-circular variance) of the tuning curves, neither of which was significantly different for PV and Pyr cells (p < 0.63 and p <0.28 respectively). In addition, we found no differences in the distribution of PDs between the two cells types (two-sample test for circular data, p=0.61) or in the tendency for tuning (X^2^ test; p=0.09; X^2^=2.17; degrees of freedom=1).

Next, we tested for correlations between the activity of individual cells and the kinematic parameters of the movements performed. Figure 4A shows the correlation throughout a trial of the activity of a PV and a pyramidal cell with movement duration (MD: the time between movement onset and when the cursor entered the peripheral target). Long before movement started, the activity of the PV (but not the pyramidal) cell was negatively correlated, on a trial-to-trial basis, with the time of the subsequent movement (Figure 4B; note that the positive correlation after movement is trivial, since this time window overlaps with the time of movement). This propensity was also present at the population level; PV cells had a stronger and earlier correlation with MD compared to the Pyr cells (Figure 4C). This was evident both at the average level of correlation and the fraction of the significant cells out of the entire population (Figure S2). Since PV and Pyr cells differed in their rate levels we tested whether cells with enhanced task-related rate modulation also expressed a stronger rate-to-MD correlation (Figure 4D). The two parameters were significantly related for PV cells (Pearson n r=-0.3; p<0.003) whereas the correlations expressed by pyramidal cells was independent of rate modulations (Pearson n r=-0.11; p<0.18).

**Figure 4:**
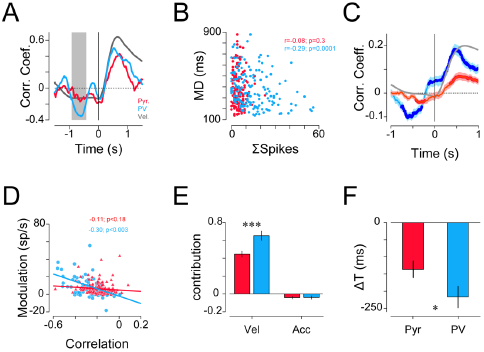
Cell-specific neural-behavioral correlation during reaching. (A) Example of a time-resolved correlation computed for two cells (red: pyramidal; blue: PV) between trial-to-trial firing rates and corresponding movement times. Correlation values were obtained by measuring single trial firing rate in a 500 ms sliding windows and correlating it with movement time. Correlation functions were aligned on movement onset. Gray box shows the time window from which the firing values were taken in panel (B). (B) Scatter plot of the movement times (y-axis) against single trial spike counts (x-axis) of each cell (red-Pyr firing and blue-PV firing) as measured for the highlighted gray box shown in A. (Pyr-MT correlation: ρ=-0.08, p=0.3; PV-MT correlation: ρ=-0.29, p=0.0001; Pearson’s correlation). (C) Average time-resolved correlation function between firing rate and movement time computed for pyramidal (red) and PV (blue) cells. Shading area corresponds to the standard error of the mean. Filled circles highlight the time points when the correlation value was significantly different from zero (Wilcoxon’s sign-rank; p<0.0006- Boneferroni’s correction for multiple comparisons- 0.01/(500/30); 500 ms is the time window and 30 ms is the time steps). Gray trace shows the average hand position (distance from the center target) recorded in parallel to the neural data. Time zero corresponds to movement onset. (D) The effect of firing rate level on firing-to-behavior correlation was estimated by plotting the relations between single cell rate modulation (relative to baseline level) measured at time point of minimal correlation and the correlation coefficient value for Pyr (red triangles) and PV (blue circles) cells. The correlation coefficient between the two parameters were computed and their significance level was estimated (Pyr: ρ=-0.11, p=0.18; PV: ρ=-0.30, p=0.003; Pearson’s correlation). The lines show the linear regression computed for each set of values (R^2^: PV=0.093; Pyr=0.012). (E) The contribution of velocity (Vel) and acceleration (Acc) to the reconstruction of firing rate (see Methods section for details. Velocity contribution: Pyr= 0.45±0.03, PV= 0.65±0.05, p<0.001; Acceleration contribution: PV=-0.04±0.02, Pyr=-0.04±0.01; Wilcoxon’s sign-rank). (I) Mean time shift (ΔT) required to achieve maximal similarity between actual neural firing and reconstructed firing (PV=-217±31.9 ms, Pyr= -134.7±24.9 ms; Wilcoxon’s sign-rank, p<0.05). See also Figure S2.

Finally, we investigated the relationship between single cell activity and movement velocity by using the instantaneous hand velocity and acceleration to build a linear model to account for the trial-to-trial firing of the recorded cells (Medina and Lisberger, 2007). Velocity made a significantly higher contribution to predicting PV firing than Pyr firing (Figure 4E). In addition, the firing-to-behavior time lag required for the best model fit for PV cells was more negative than for the Pyr cells (Figure 4F), consistent with our previous finding that PV task-related firing preceded that of Pyr cells.

Taken together, these results suggest that the movement-related activity of PV cells starts before that of excitatory cells, but nevertheless robustly encodes motor parameters such as direction, movement duration and velocity. This finding assigns a dominant role to PV neurons in coordinating not only the activity of pyramidal neurons, but also in steering motor behavior.

### Synchronous activation of PV and Pyr cells

SCP-responsive PV and Pyr cells are not independent channels but rather embedded within a rich cortical network which makes them likely to interact. These interactions can amplify or suppress incoming cerebellar signals in a task-dependent manner. We tested this possibility using two different approaches. First, we estimated the correlated firing (spike-count correlation) of simultaneously recorded PV and Pyr cells triggered by SCP stimulation. This analysis was only carried out on cells recorded simultaneously from two different electrodes, to prevent any masking of synchrony due to super-positioning of action potentials (Bar-Gad et al., 2001). We found that SCP stimulation triggered cell-specific synchronization patterns. Pairs of PV and pyramidal cells expressed an early increase in correlated firing followed by a decrease in co-firing which occurred at time lags consistent with PV-to-Pyr inhibition (Figure 5A). This correlation pattern is consistent with the expected correlation calculated for a simulated TC circuit with FFI (Figure 5B-C and Figure S3) constructed based on previously published parameters (Cruikshank et al., 2007; Izhikevich, 2003; Kremkow et al., 2010). By contrast, the synchronizing effect of SCP stimulation on pairs of pyramidal cells was substantially weaker (Figure 5D). Here again, the results were consistent with the correlation calculated for the simulated Pyr-Pyr pairs using similar conditions to our stimulation protocol in terms of number of sweeps (Figures 5E-F).

**Figure 5:**
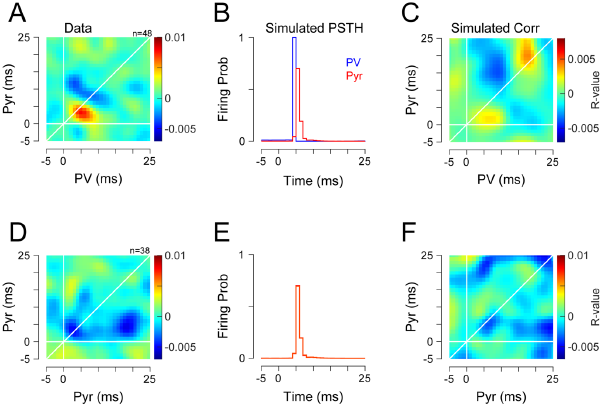
Joint Peri-stimulus time histogram of PV and pyramidal pairs. (A) Average Joint peri-stimulus time histogram (jPSTH) of simultaneously recorded pairs of PV and pyramidal cells (n=48). Time zero is stimulation time. Color scale corresponds to the normalized correlation values (actual counts – expected counts over the standard deviations). Vertical and horizontal white lines indicate the time of the stimulation. Diagonal line corresponds to the zero-lag time between the two cells. (B) “Stimulus-triggered response” of simulated cell pairs (n=100 pairs; 400 trials) as obtained from a model of a feed-forward inhibition circuit. Responses were aligned on thalamic spike times (t=0). (C) Average jPSTH computed between simulated pairs of PV and Pyramidal cells that resulted from the simulated circuitry as shown in B. (D-F) same as A-C but for pairs of pyramidal cells. The number of recorded pairs used to compute (D) was n=38 and for F number of simulated pairs was the same as in C (n=100 pairs; 400 trials). See also Figure S3.

Next, we searched for traces of this SCP-triggered correlated firing in the activity of the cells during task performance. During natural behavior the activation of the CTC pathway is sparse and dispersed compared to the tightly synchronized activation in response to SCP stimulation. Nonetheless, we found that around movement onset, cells that are part of the CTC system were co-driven in a cell-specific manner consistent with their activation pattern in response to SCP stimulation. Specifically, pyramidal and PV cells expressed time-varying synchrony locked to movement onset (Figure 6A), whereas pairs of Pyr cells were generally and consistently uncorrelated (Figure 6B). The correlation pattern between PV and Pyr cells consisted of a positive correlation before onset and after the offset of movement and a decorrelation during movement (Figure 6C; p<0.02), whereas the correlation of Pyr cells was not significantly modulated between these epochs (Figure 6C; p=0.72). The observed correlation for pairs of PV/PV and Pyr/PV were similar in pattern to that found in response to evoked CTC activation, whereas the Pyr/Pyr correlation was substantially weaker. Similar results were obtained when measuring the correlated firing only in trials directed towards the preferred direction (Figure S4). Finally, to test the effect of the different rate levels of PV and Pyr cells, we computed the correlated firing between a subset of Pyr-Pyr pairs where at least one of the neurons had a rate level comparable (i.e., equal to or higher than the mean-½SD) to that of the PV cells (Figures 6D-F). We found that although this subset of pyramidal cells exhibited a similar response amplitude around movement onset as the PV population (Figure 6D), the cells expressed no correlation around movement onset (Figure 6E). In addition, the relationship between the task-related firing of cells and the corresponding level of correlated firing was not significant (Figure 6F). Taken together, these results suggest that in general, the CTC drive is sufficiently strong to induce PV-to-Pyr correlated firing around movement onset, so that the interactions between these cells are further amplified and may thus affect subsequent movement.

**Figure 6:**
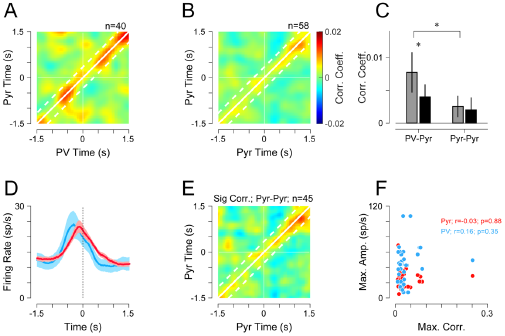
Joint Peri-event time histogram of PV and Pyramidal pairs around movement onset. (A) Average Joint Peri-event time histogram (jPETH) for pairs of PV and pyramidal neurons around movement onset. White horizontal and vertical lines denotes movement onset time (t=0). Solid diagonal line corresponds to zero inter-unit lag time while dashed lines corresponds to the time around the diagonal within which the average was calculated for panel C. Color scale corresponds to the R-value of the correlation martix ((counts-predicted)/std; n=40 pairs). (B) Same as (A) for pyramidal-pyramidal pairs (n=58 pairs). (C) The average correlation at baseline (pre-movement time, gray bars) and during movement (black bars), was calculated during a time window spanning -600 to -100 ms relative to movement onset and -100 to +400 ms around the same event. The inter-unit time lags that were considered for computing the averaged correlation were ±350 ms (shown as diagonal lines in panels A and B). Each bar represented the mean correlation value computed across pairs and the corresponding s.e.m. (PV-Pyr pairs; n=40 pairs; Base: 0.008±0.003; Move: 0.004±0.002; Wilcoxon’s sign-rank test; p<0.02; Pyr-Pyr pairs; n=58 pairs; Base: 0.0025±0.001; Move: 0.002±0.0019; p=0.72). During pre-movement period (baseline) the baseline correlation between PV-Pyr and Pyr-Pyr pairs was significantly different (p<0.02). (D) The average movement related activity of PV and a subset of Pyramidal neurons that had a specifically high response amplitude around movement onset (n=45 neurons). Responses were aligned and movement onset time and shaded areas show the s.e.m. (E) Same as B, but for computed for the subset of Pyramidal cells shown in D. (F) Relationship between maximal, zero-lag correlation found for each Pyr-to-PV cell pairs and the maximal firing rate around movement onset exhibited by PV (blue circles) or Pyr (red) neurons (Pyr: ρ=-0.03, p=0.88; PV: ρ=0.16, p=0.35; Pearson’s correlation). See also Figure S4.

## DISCUSSION

Motor cortical circuitry integrates multiple inputs and transforms them into descending motor commands. Of these inputs, cerebellar signals are important for motor timing and coordination (Holmes, 1917, 1939; Horne and Butler, 1995). This supposition relies on data obtained from individuals suffering from cerebellar lesions or injury. However, it is unclear how cerebellar outflow, which is funneled through a relatively small anatomical pathway, regulates the timing of motor cortical activity and hence motor actions. This study describes a novel effort to address this question by combining electrophysiological and pharmacological tools to dissect the motor cortical circuit that integrates cerebellar signals in behaving primates. We showed that the CTC system follows the same organizational principles as reported for sensory systems; namely, feedforward inhibition through enhanced activation of PV cells compared to pyramidal cells. We further showed that the functional outcome of this arrangement is the earlier onset and tighter relationship to movement duration of the activity of PV cells compared to pyramidal cells. We therefore suggest that the dual, inhibitory-excitatory activation of motor cortical cells by CTC input is important for prioritizing cerebellar signals compared to competing inputs, and that this mechanism enables the CTC system to play a dominant role in timing movement onset. This arrangement assigns a functional role to the potent cerebellar link with the motor cortical inhibitory system.

### Combining electrical and pharmacological perturbations to disentangle motor cortical circuitry

The lack of information about cell type and its input/output connectivity pattern constitute major obstacles to interpreting the functional consequences of single cell activity recorded in behaving primates. Here we combined two approaches to overcome this limitation. We used SCP stimulation to identify motor cortical cells which integrate cerebellar signals (Nashef et al., 2018a). We then pharmacologically identified PV and pyramidal cells in this population. To do so, we relied on data previously obtained for thalamocortical systems in rodents which showed that thalamic inputs synapse almost exclusively onto pyramidal and PV cells (Hull et al., 2009; Ji et al., 2015) and that these synapses are mediated by GluR2 lacking AMPA receptors on the PV but not the pyramidal cells (Geiger et al., 1995; Jonas et al., 1994; Kondo et al., 1997; McBain and Dingledine, 1993). We classified cells for which the early SCP-triggered response was abolished in the presence of NASPM, an antagonist of GluR2- lacking AMPA receptors, as inhibitory PV interneurons. A possible confound to our interpretation of the NASPM results is whether the TC projection to inhibitory interneurons in the primate motor cortex primarily targets PV cells, as is the case in sensory systems in rodents, since their cortical organization and layer-specific thalamic input are different. The response properties of the interneurons identified here (i.e., higher baseline firing rates, narrow action potentials and monosyantpic connectivity with TC fibers) are all supportive of these cells being PV interneurons. Nonetheless, our functional conclusions are not dependent on the exact subtype of inhibitory cell receiving TC input.

Another possible confound is the non-specific and varied effects of polyamines on NMDA receptors, which depend on the concentration and specific chemical subtype (Rock and Macdonald, 1995). Although NMDA receptors are not an integral part of the FFI circuity (Hull et al., 2009; Kloc and Maffei, 2014), they may play an indirect role in modulating the response pattern of cortical cells to SCP activation, especially the late inhibitory phase. However, NASPM was found to not affect NMDA-mediated activation (Sanderson et al., 2016). Moreover, our analysis of responses to SCP stimulation was restricted to the early excitatory response, which is not affected by NMDA-mediated currents. We therefore conclude that the NASPM-induced attenuation of SCP-triggered responses were primarily mediated by GluR2- lacking AMPA receptors and can thus be used to accurately identify PV cells.

We found that AP width reliably distinguished between PV and pyramidal neurons identified by SCP-stimulation and NASPM responses. This criterion is commonly used to distinguish between excitatory and inhibitory cells both in rodents (Bortone et al., 2014; Middleton et al., 2012; Zagha et al., 2015) and in primates (Kaufman et al., 2010; Kaufman et al., 2013; Merchant et al., 2008; Mitchell et al., 2007; Onorato et al., 2019). However, recent findings indicate that pyramidal tract neurons (which are exclusively excitatory cells) can have a narrow AP which is related the existence of Kv3.1b potassium channels (Soares et al., 2017). Our results may not necessarily be at odds with this finding, since our dataset was restricted to SCP-responsive cells; i.e., cells that receive CTC input, and the fraction of PT neurons within this population is unclear (Peters et al., 2017).

### Feedforward inhibition (FFI) as a canonical TC mechanism across species and modalities

We found a large fraction of cells in the motor cortex that received TC input via NASPM-sensitive synapses, indicating inhibitory cells (a ratio of 1-to-2 between SCP-responsive PV to pyramidal cells). The existence of a direct connection between the CTC system and inhibitory interneurons indicates that cerebellar outflow triggers FFI in the motor cortex of behaving primates, similar to what has been observed in sensory and frontal cortical areas of rodents (Cruikshank et al., 2007; Delevich et al., 2015; Gabernet et al., 2005; Swadlow, 2003). The fundamental principle of FFI is that driving signals simultaneously trigger excitation and inhibition in the target structure. The functional benefits of this arrangement are still not fully understood. Several studies have speculated that it may provide a temporally-tight perceptual window within which sensory information can be integrated (Bruno, 2011). It was further suggested that this arrangement makes cortical neurons sensitive to synchronized thalamic input (Bruno, 2011; Bruno and Sakmann, 2006; Wang et al., 2010). In the motor pathways, the same FFI architecture can thus have similar effects on single cell activity but very different behavioral consequences. We previously posited that the excitation/inhibition interplay triggered by the CTC input plays a role in generating the firing transient at the onset of movement (Nashef et al., 2018a). Consistent with this hypothesis, blocking the cerebellar outflow by dentate cooling (Hore and Flament, 1988) or high frequency stimulation in the SCP (Nashef et al., 2019) abolishes this transient, particularly in cortical cells that integrate CTC input. The loss of this transient was accompanied by ataxic movements. These results suggest that thalamo-cortical FFI may play essentially the same role in the temporal organization of activity in both the motor and sensory areas.

The direct and potent contact of inhibitory cells by the CTC system may have additional consequences beyond the temporal organization of cortical activity. Non-invasive cerebellar stimulation in humans (using TMS or tDCS) produces a general inhibition of M1 termed CBI (Ugawa et al., 1991; Ugawa et al., 1995), which is correlated with motor learning (Schlerf et al., 2012). Since the CTC is an excitatory pathway, CBI has been attributed to activation of inhibitory Purkinje cells (Celnik, 2015). Our findings may revise and extend this hypothesis by indicating that CBI could be related, at least in part, to the CTC activation of motor cortical inhibitory interneurons.

Calcium-permeable AMPA receptors makes TC synapses not only particularly strong, but also prone to plastic changes (Isaac et al., 2007; Plant et al., 2006). This could explain the fact that during motor learning TC input is often modified (Biane et al., 2016), and that the CBI is correlated with motor learning. Interestingly, the concept of learning via changes in inhibitory connections has been supported by recent theoretical models (Haga and Fukai, 2019; Luz and Shamir, 2012; Mongillo et al., 2018; Vogels et al., 2011).

### PV interneuron steer cortical activity and motor output

Cortical PV interneurons are commonly assigned different functional roles. In sensory areas, PV cells are implicated in shaping the tuning of nearby pyramidal cell responses (Atallah et al., 2012; Hofer et al., 2011; Kerlin et al., 2010; Merchant et al., 2008; Sohya et al., 2007). These cells have also been implicated in maintaining the excitation-inhibition balance needed to regulate cortical activity (Isomura et al., 2009; Xue et al., 2014). Our findings suggest that the PV cells of the motor cortex play a specific role in the temporal organization of neural activity and hence motor behavior. We found that around movement onset, PV cells are recruited before pyramidal cells, probably due to the stronger input PV cells receive from the CTC system. Similar findings were reported for PV neurons in sensory areas (Yu et al., 2019) and in the motor cortex of rodents making spontaneous reaches (Estebanez et al., 2017). However, the time lag between response onset for the two cells types reported here (~50 ms) was much larger than in the somatosensory cortex (a few ms). This discrepancy could reflect different properties of both the thalamic volley (e.g., more slowly ramping thalamic activity could produce a larger PV-to-pyramidal time lag) and the cell properties of the PV neurons in different cortical areas (Tasic et al., 2018), such as TC synaptic efficacy, threshold for activation, etc. From a functional perspective, the short PV-to-Pyr time lag found in sensory areas might be optimal for imposing a fast perceptual time window (Bruno, 2011), whereas the longer time-lag found in the motor cortex is more appropriate for timing movements that involve longer time scales.

The functional implications of this early onset of inhibitory signals remain unclear. It is possible that the temporal order of events, together with the enhanced PV-to-Pyr synchrony of SCP-responsive cells, are important for the appropriate onset of movement. At any given point in time, cortical cells integrate a multitude of converging inputs, not all of which are related to the ensuing movement. Thus, early inhibition could suppress these inputs and allow for the subsequent, movement-related TC excitation to more effectively recruit downstream elements. In this manner, the gain of the thalamic input is enhanced, despite its modest synaptic weight. If true, this arrangement predicts that in the absence of CTC drive, task-related activity of PV and pyramidal cells will overlap in time, reaction time will increase (since it will take more time and effort for activity to produce movement) and the neural and behavioral variability at movement onset will increase (since the initial suppressive effect of PV neurons will not be available).

The early onset of PV cells puts them in an excellent position to steer movements, but the finding that these cells are more strongly correlated with kinematic parameters (i.e., velocity) seems counterintuitive, since pyramidal cells are the output stage of the motor cortex that transmit signals to downstream elements. The stronger synaptic input to PV neurons may make them more easily driven by thalamic input; hence, task-related activity may be more homogeneous across the population of PV cells. In contrast, pyramidal cells are more diverse and their activity is composed of a mixture of multiple (high- and low-level) parameters. As such, although the population activity of pyramidal cells may be strongly correlated with kinematic parameters and drive movements, this relationship is weaker for the activity of single Pyr cells than for single PV cells.

### Summary

We report a novel dissection of motor cortical circuitry that integrates cerebellar input during motor behavior. By combining electrophysiological and pharmacological tools, we identified PV and pyramidal neurons in the motor cortex of behaving monkeys. We showed that the two groups of cells have different firing properties (consistent with data from previous studies) but also different patterns of task-related activity. The data indicate that thalamic input in the motor cortex of behaving primates triggers feedforward inhibition, similar to observations in sensory systems. In parallel, we found that PV cells fire earlier than pyramidal cells at movement onset, consistent with the dominant role of cerebellar signals (mediated through the thalamus) at this time. In addition, we found that PV cells are related to task parameters in a similar or stronger manner than pyramidal cells. We therefore suggest that PV neurons are important for prioritizing cerebellar signals at movement onset through the early suppression of competing uncorrelated firing. In this way, these cells act to improve the signal-to-noise ratio for thalamic input, which otherwise might be too weak to properly activate the relevant effectors. The strong thalamic drive to PV cells makes them a more coherent group of neurons (compared to pyramidal cells) and enhances their correlation with kinematic parameters as compared to pyramidal cells. Finally, we suggest that although the operational principle of FFI in general and PV neurons in particular is similar across cortical regions and species, the temporal properties of the motor cortical circuity are tuned to controlling movement timing, rather than integrating percepts as in the sensory cortices.

## ACKNOWLEDGEMENTS

This work was funded by the Israel Science Foundation (ISF-1801/18) the Jerusalem Brain Center (to AN) and the generous support of the Baruch Foundation (to YP).

## CONTRIBUTION

YP and SIP conceived the study. AN performed the experiments and analyzed the data, YP, AN and SIP wrote the manuscript.

## DECLARATION OF INTEREST

The authors declare no competing interest.

## STAR METHODS

### RESOURCE TABLE

| Reagent or Resource | Source | Identifier |
| --- | --- | --- |
| Experimental model: Organisms/strains |  |  |
| *Macaca fascicularis* | The Hebrew University | N/A |
| Software and Algorithms |  |  |
| MATLAB 2016b | Mathworks | https://www.mathworks.com/ |
| Drugs |  |  |
| 1-Naphthylacetyl spermine trihydrochloride | Sigma Aldrich | https://www.sigmaaldrich.com/united-states.html |
| Other |  |  |
| NSEX-100 | NSEX-100 | NSEX-100 |

## CONTACT FOR RESOURCE SHARING

Further information and requests for resources and reagents should be directed to and will be fulfilled by the lead contact, Yifat Prut.

## EXPERIMENTAL MODEL AND SUBJECT DETAILS

This study was performed on three adult female monkeys (*Macaca fascicularis*, weight 4.5- 8 kg). The care and surgical procedures of the subjects were in accordance with the Hebrew University Guidelines for the Use and Care of Laboratory Animals in Research, supervised by the Institutional Committee for Animal Care and Use.

## METHOD DETAILS

### Behavioral task

Data were obtained from three Macaca fascicularis monkeys (females, 4.5-8 Kg). Primate care and surgical procedures were in accordance with the Hebrew University Guidelines for the Use and Care of Laboratory Animals in Research, supervised by the Institutional Committee for Animal Care and Use. The behavioral task used here was described in our previous study (Nashef et al., 2019). Briefly, the three monkeys were trained to sit in a primate chair, wear a unilateral exoskeleton (KINARM, BKIN technologies), and perform an arm reaching task. The exoskeleton constrained the arm to move in a horizontal plane at the level of the shoulder, and allowed free movement about the elbow and shoulder joints in that plane. The end point coordinates of the exoskeleton (i.e., the monkey’s hand position) controlled the location of a cursor on a horizontally positioned screen in front of the animal (Figure 1A). In this task, the monkeys were instructed to move the arm to locate a cursor within a central target. After 500 ms, a peripheral target (one of 8 evenly distributed targets) appeared. After a fixed delay time of 300-750 ms the central target disappeared (“GO” signal), cuing the monkey to reach to the visible peripheral target. If the monkey moved the cursor to the correct target within the predefined time limits it was rewarded with a drop of applesauce. To encourage the monkey to predict the timing of the “go” signal, we limited the total time it had to reach the peripheral target after the GO signal to 500 ms and inserted a 200 ms grace period before the GO signal. Onset of movement within this time frame did not abort the trial.

After training was completed, a recording chamber (21×21 mm) was attached to the monkeys’ skull above the hand-related area of the motor cortex in a surgical procedure under general anesthesia. After a recovery and re-training period, we recorded motor cortical activity extracellularly by advancing electrodes through the chamber in daily recording sessions.

### Insertion of stimulating electrode into the superior cerebellar peduncle (SCP)

To insert a chronic stimulating electrode into the SCP ipsilateral to the trained arm, we implanted a small chamber above the cerebellum (based on a macaque neuro-atlas) and used a post-surgery MRI to plan an electrode trajectory to the SCP. A bi-polar concentric electrode (NSEX100, David Kopf Instruments, impedance range of 30-60 k?) was positioned in the peduncle through the chamber. Evoked intra-cortical responses to stimulation through the electrode were used to verify its location before gluing the electrode to the chamber (Nashef et al., 2019; Nashef et al., 2018a; Nashef et al., 2018b; Ruach et al., 2015).

### SCP stimulation protocol

Biphasic (200 μs each phase) stimulation pulses were applied to the SCP electrode while the monkey performed the task and cortical activity was recorded. A single set of stimuli consisted of about 200 stimuli that were delivered at 3 Hz at a fixed intensity (ranging from 50 to 300 μA).

### Recording

Recordings were obtained using one of two methods:

1- *Iontophoresis sessions*: For a full description of the method, see (Thiele et al., 2006). During the iontophoresis sessions, three-barrel pipettes were inserted through the chamber to different cortical sites, mostly in the primary motor cortex (M1) which was defined based on MRI scans, their relation to anatomical landmarks (i.e., the central and arcuate sulci) and threshold for motor response. The central barrel in the pipette contained a tungsten electrode (impedance 300-800 kΩ at 1,000 Hz) for extracellular recording. Before each recording session, the two peripheral barrels were filled with NASPM (or glutamate) using polyamide hypodermal injection tubing (purchased from World Precision Instruments, Ltd.), and a bare tungsten wire was inserted into each barrel and held in place using a drop of glue. The signal obtained from each electrode was amplified (×10^4^), bandpass-filtered (300-6,000 Hz), digitized (32 kHz), and saved to disk. NASPM and glutamate were ejected from the pipette by applying a positive or negative current, respectively, to the tungsten wires (30-200 nA). A -15 nA retention current of the opposite sign was applied to the wires at other times.

2- *Other sessions*: Glass coated tungsten electrodes (impedance 300-800 kΩ at 1,000 Hz) were inserted through the chamber to different cortical sites, mostly in M1. The signal obtained from each electrode was amplified (×10^4^), bandpass-filtered (300-6,000 Hz), digitized (32 kHz), and saved to disk. Recordings were made with up to 4 individually moveable electrodes (Flex-MT by Alpha Omega, Nazareth, Israel).

### NASPM preparation

1-Naphthylacetyl spermine trihydrochloride (NASPM, Sigma-Aldrich; molecular weight= 479.91 g/mol) was used for blocking GluR2-lacking AMPA mediated receptors on parvalbumin (PV) cells. Preparation of NASPM started with dissolving the powder (5 mg) into 2cc of saline (concentration 5mM). The solution was kept at -80 C^o^ and warmed before use. For injection, we diluted the solution further with saline to obtain a final concentration of 1mM and pH of 5-6; the final solution was not kept for more than 7 days after preparation. We recorded responses of cortical neurons to glutamate iontophoresis in one session per monkey for control. Here, 100 mg of L-glutamate was dissolved in 6.796 ml of artificial CSF and titrated with HCO_3_^-^ to increase the pH to between 7.4-7.8.

## QUANTIFICATION AND STATISTICAL ANALYSIS

### Cell classification

We combined naïve Bayesian and support-vector machine (SVM)(Cortes and Vapnik, 1995) classifiers to identify PV and pyramidal cells. First, we trained the classifier with data from cells that were tested using NASPM (n=58). For each cell we calculated several features to guide the classification: width of action potential, amplitude of waveform (peak-trough), slope of waveform (amplitude/width), decay of waveform after peak, peak-to-trough (peak/trough), amplitude to SCP stimulation, gain after SCP stimulation, area of excitatory response following SCP stimulation, area of inhibitory response following SCP stimulation and pre-stimulus firing. This step was carried out to find the parameters that would yield maximal performance. We assessed the performance of the classifier for all the combinations of the 9 features mentioned (overall, 511 different combinations). For each combination a relevant-component analysis (RCA) was performed over the data, to optimize the distance between the two cell classes(Bar-Hillel et al., 2003; Shental et al., 2002). Next, we ran the classifier 100 times for each of the cells in the training set (n=58), where at each iteration a random 60% selection of the remaining cells (n=34) were considered. In practice this means that in each iteration the classifier was retrained with the randomly selected subset of 34 cells in order to minimize the effect of noisy labels. After the 100 iterations, the most common label in both classifiers was considered to be true. For instance, if the naïve Bayesian labeled 70 cases (out of 100 iterations) as PV and SVM labeled 55 cases as Pyr, the naïve Bayesian result, PV, was taken as the final classification. We found that width, pre-stimulus firing activity and excitation following SCP stimulation were the most important features in distinguishing between the two subpopulations providing an accuracy of 75% in classification (76% of the PV and 75% of Pyr), based on the cells tested with NASPM. The classifier was then applied to the cells that were not recorded during NASPM application (the unknown labels), in the same way.

### Data analysis

#### Stimulus-evoked responses of single neurons

We identified SCP-responsive cortical neurons according to a previously published method (Nashef et al., 2019; Nashef et al., 2018a).

#### Simulation of a feed-forward inhibition (FFI) circuit

We simulated an FFI circuit using a previously published (Izhikevich, 2003) excitatory\inhibitory (E\I) cortical network (http://izhikevich.org/publications/spikes.htm) with randomized incoming thalamocortical input. To mimic the SCP stimulation, we enhanced the randomized input at time zero. Furthermore, we used parameters that were found in the past for the strength of thalamocortical input on PV and pyramidal neurons (Cruikshank et al., 2007), specifically, the conductance of the thalamic-PV synapse was set to 3 nS and thalamic-Pyr. to 0.7 nS, to ensure reliable inhibition in our circuitry. Other well defined parameters were: spiking threshold=-57 mV; resting potential= -70 mV; τ(Pyr)=2 ms; τ(PV)=10 ms. The intracortical excitatory synapse conductance at 1 nS and the inhibitory at 2 nS (Kremkow et al., 2010). To mimic our experiment, we simulated 400 trials of incoming input for each connected pair, averaged the firing probability and found the jPSTH for these “simultaneous” 400 trials. To insert noise in the response, at each trial we randomized the connectivity strength between the PV cell and the pyramidal cell, and also the auto-connectivity of the individual cell. At the end, for each pair we had two (one PV and one Pyr) matrices of 400 x time, where each row is the firing of the cells around one CTC input. This whole process was performed 50 times, to resemble different pairs and then the average response and average jPSTH were found for these 50 pairs, using bin-by-bin averaging.

#### Task-related response properties of recorded neurons

##### 1- Background firing

The background firing of each neuron was calculated as the average firing rate when no task-related modulation occurred, in the time window between 2 and 1 seconds before movement onset (−2 to -1 sec).

##### 2- Preferred Direction

For each single unit, we computed the tuning function and its preferred direction (PD). The preferred direction was calculated individually for each isolated unit using a resampling method (Shalit et al., 2012) (4,000 repetitions) during the -500 to 500 ms around movement onset.

##### 3- Orientation selectivity index (OSI)

First we calculated the tuning curve of each cell, and then the OSI was defined as 1 – *r*, where *r* is the circular variance of the tuning curve (Atallah et al., 2012).

##### 4- Onset time of the task-related activity

To calculate the response onset time of neurons while overcoming their noisy background firing we applied the following steps. First, time of movement onset was defined using the XY-position signals that are continuously recorded in the KINARM setup. A dedicated algorithm identified transitions in the distance of the instantaneous XY position of the cursor to the center of the central target (Asher et al., 2010). Next, we calculated the average peri-event time histogram (PETH, background firing subtracted) for each neuron around movement onset and found the maximal firing rate in the window ±1 sec around movement onset. We then normalized the firing rate by the maximum rate and, starting at the peak time we moved backwards in time until the point when the firing rate dropped below 0.2 (20% of the peak) for the first time. This was defined as the onset of the movement-related firing.

##### Joint Peri-Stimulus Time Histogram

We based our quantification of the joint PSTH on previous work (Aertsen et al., 1989). First, we used the PSTH’s of two neurons to calculate the raw JPSTH matrix, in which the *(i,j)-th* bin was the time of coincidence in which one neuron spiked at the *i^th^* bin and the second spiked at the *j^th^* bin. To correct for rate modulation we calculated the PSTH predictor (Aertsen et al., 1989). The predictor matrix is the product of the single neuron PSTHs; i.e., the *(i,j)-th* bin is equal to PSTH1*(i)*\*PSTH2*(j)*. The final jPSTH was calculated as: jPSTHPredict – jPSTH_Raw_. The jPSTH was calculated in bins of 1 ms and smoothed with a two-dimensional Gaussian window with a SD of 1.5 ms.

##### Neural-Behavioral correlation

We quantified the relationships between neural firing and movement duration, MD by measuring for each cell the time resolved correlation between the vector of spike counts (counted for each trial in a given time window) and the MD found for those trials. For this analysis we only considered cells with at least 20 available trials and a significant tuning. To compute the correlations between spike counts and MD, available trials were first aligned on movement onset (from -1500 to +1500 ms around this event) and spike numbers were counted for time windows of 200 ms, advancing in steps of 10 ms. For each window, we computed the correlation coefficient between spike counts and MD. Correlation coefficient were then averaged across all cells to obtain the population-based correlation between rate and MD. The standard error of the mean was then computed per bin for each class of cells.

## SUPPLEMENTARY FIGURES

**Figure S1:**
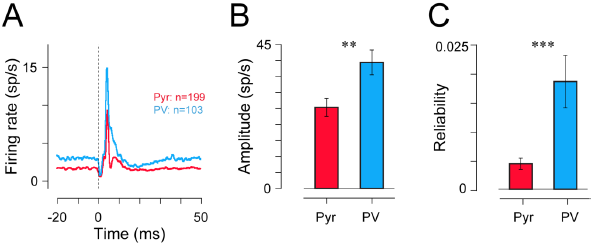
SCP response properties of classified cells, related to Figure 2. (A) The average response to SCP stimulation of classified Pyr and PV cells. Vertical line corresponds to the stimulation time (Pyr- n=199 cells and PV- n=103 cells). (B) Comparison of the amplitude of the different classes of cells following SCP stimulation, measured as the maximal firing rate following the stimulation (Pyr: 25.3±2.8; PV: 39.4 ±3.9 sp/s; Wilcoxon’s sign-rank; p<0.005). (C) Comparison of the response reliability (Wang et al., 2010) of the different classes of cells following SCP stimulation (Pyr: 0.004±0.0004; PV: 0.019±0.003; Wilcoxon’s sign-rank; p<0.001).

**Figure S2:**
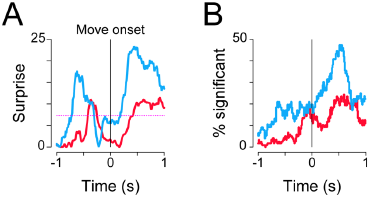
Cell-specific neural-behavioral correlation during reaching, related to Figure 4. (A) Mean surprise values (-log(p)) computed for the correlation functions shown in (Figure 4C), dashed line denotes the threshold for determining significance (i.e., surprise value of 7.4). (B) Fraction of cells that exhibited significant correlation between rate and movement time as computed in Figure 4C.

**Figure S3:**
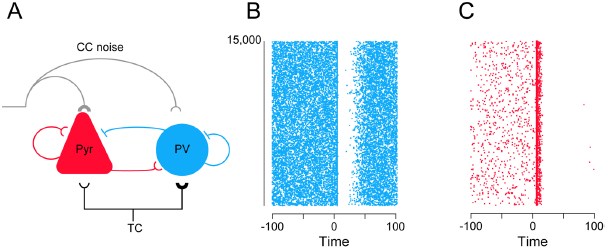
Simulated Feed Forward Inhibition circuitry, related to Figure 5. (A) The synaptic connections that were used to model the feed-forward inhibitory circuity. Both PV and Pyr cells received an excitatory input from thalamocortical neurons (TC)- the input to PV was however 4 times stronger than the input to Pyr neurons. Each cell then contacts the other cell and itself (autoactivation for Pyr and autoinhibition for PV cell). Both cell types also receive a corticocortical noisy input. (B) Raster of PV response to simulated TC input. Each line corresponds to the PV response to one TC spike, and time zero is the TC spike time. (C) Same as B, but for Pyr cell.

**Figure S4:**
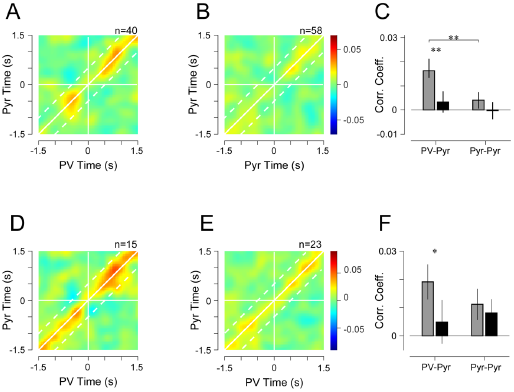
Joint Peri-event time histogram of PV and Pyramidal pairs around movement onset, related to Figure 6. (A) Average jPETH for pairs of PV and pyramidal neurons around movement onset. White horizontal and vertical lines denotes movement onset time (t=0). Solid diagonal line corresponds to zero inter-unit lag time while dashed lines corresponds to the time around the diagonal within which the average was calculated for panel C. Color scale corresponds to the R-value of the correlation martix (counts-predicted)/std; n=40 pairs). The jPSTH was aligned around the PD of one unit of the pair randomly. (B) Same as (A) for pyramidal-pyramidal pairs (n=58 pairs). (C) The average correlation at baseline (pre-movement time, gray bars) and during movement (black bars), was calculated during a time window spanning -600 to -100 ms relative to movement onset and -100 to +400 ms around the same event. The inter-unit time lags that were considered for computing the averaged correlation were ±350 ms (shown as diagonal lines in panels A and B). Each bar represented the mean correlation value computed across pairs and the corresponding s.e.m. (PV-Pyr pairs; n=40 pairs; Base: 0.017±0.004; Move: 0.003±0.004; Wilcoxon’s sign-rank test; p<0.006; Pyr-Pyr pairs; n=58 pairs; Base: 0.004±0.003; Move: -0.0003±0.004; p=0.29). During pre-movement period (baseline) the baseline correlation between PV-Pyr and Pyr-Pyr pairs was significantly different (p<0.01). (D) Same as (A) but for pairs of units with similar PD- pairs that have similar or neighboring PD (n=15). (E) Same as (A) for pyramidal-pyramidal pairs, that have similar PD (n=23). (F) Same as (C), for pairs of cells with similar PD (PV-Pyr pairs; n=15 pairs; Base: 0.019±0.006; Move: 0.005±0.007; Wilcoxon’s sign-rank test; p<0.05; Pyr-Pyr pairs; n=23 pairs; Base: 0.01±0.006; Move: 0.008±0.005; p=0. 95).

## Notes

### Competing Interest Statement

The authors have declared no competing interest.

